# Single-cell transcriptome analysis reveals evolutionarily conserved features during the transition from normal breast stromal cells to cancer-associated fibroblasts

**DOI:** 10.1101/2022.05.05.490693

**Authors:** Ana Paula Delgado, Alice Nemajerova, Brian J. Nelson, Manisha Rao, Jinyu Li, Natalia Marchenko, Jonathan Preall, Ute M. Moll, Mikala Egeblad, Scott Powers

**Affiliations:** Department of Pathology and Cancer Center, Renaissance School of Medicine, Stony Brook, New York; Graduate Program in Genetics, Stony Brook University, Stony Brook, New York; Department of Applied Mathematics and Statistics, Stony Brook University, Stony Brook, New York; Cold Spring Harbor Laboratory, Cold Spring Harbor, New York

## Abstract

To comprehend cancer-associated fibroblasts (CAFs) origins, single-cell RNA sequencing was conducted on normal and cancerous breast tissue from mice and humans. We found three conserved CAF subtypes, which based on GOterm analysis we designated as matrix CAFs-, chemokine CAFs, and contractile CAFs. Matrix and chemokine CAFs originated from resident fibroblasts, while contractile CAFs originated from normal pericytes. Both human and mouse CAFs displayed upregulated genes involved in extracellular matrix organization, cellular respiration, and cell migration. Key transcription factors in both species included NFKB1, SP1, TP53, and TWIST2. Trajectory inference suggested that in some cases a transitory state characterized by JUN expression precedes the maturation of CAFs. Computational analysis revealed a common mechanism for CAF education involving the overexpression of TGF-β, PDGF, TNF, and NOTCH-family ligands in different tumor microenvironment cell types, along with reciprocal overexpression of receptors in CAFs. These findings bolster and broaden current understandings of CAF genesis.

## Introduction

Although there are a variety of theories on the origin and development of CAFs, most studies indicate that CAFs develop from tissue-resident mesenchymal cells^1–3^. Studies using human tissue have shown that there are progressive changes in fibroblastic stroma from early pre-malignant lesions onwards towards invasive carcinoma^4^. How these progressive changes are orchestrated is not completely understood, but they closely resemble changes in fibroblasts during wound healing. During both wound healing and tumor formation, tissue-resident mesenchymal cells multiply and change cell shape and other processes by stimulation with growth factors, including transforming growth factor-β (TGFβ), platelet-derived growth factor (PDGF), as well as cytokines such as IL-1 or IL-6^2,3^. Many of these factors are produced by tumor epithelial cells, but tumor-associated immune cells are known to produce them as well^2,3^.

Although single-cell RNA-sequencing (scRNA-seq) technologies in principle can expand our knowledge of CAF origins and the mechanisms by which they develop, to date single-cell RNA-sequencing CAF studies in breast cancer have been largely focused on the characterization of the complexity and heterogeneity of CAF subpopulations^5–7^. Our principal objectives in this study were to acquire novel insights into CAF development and to assess the degree to which established features of CAF development are correct. Our approach was to compare single-cell transcriptomes of normal mammary fibroblasts to CAFs in both humans and mouse models. In addition to enrichment analysis, we performed trajectory analysis of expression changes in CAF development using Monocle3 and scFates^8,9^. Additionally, we performed analysis of the microenvironmental circuits driving CAF education using Liana and NicheNet ^10^ .

## Results

### Three transcriptional subtypes of CAFs in murine PyMT breast tumors

We conducted scRNA-seq on mammary tumors from four MMTV-PyMT mice. Utilizing Seurat, we identified ten distinct clusters, including four tumor epithelial cell clusters expressing *Epcam*. These clusters varied in expression of differentiation and proliferative markers. Notably, the proliferative cluster was labeled as *Birc5*, the differentiated cluster as *Scgb2b27*, the stem-like cluster as *Sox9*, and the myoepithelial cluster as *Krt14*. Additionally, we detected two Ptprc+ immune cell clusters, one comprising lymphocytes (*Nkg7*) and the other containing myeloid cells (*Cd74*). There were four stromal cell clusters, including *Pecam1*^+^ endothelial cells and three different types of CAFs, labeled based on their selective expression of either *Col1a1*, *Col3a1*, or *Col4a1* (Figure 1A). GO term enrichment analysis of genes selectively expressed in the each of the CAF clusters compared to all other cells within the tumor indicated that the three CAFs have selective biological functions: extracellular matrix organization, chemokine activity, or contraction (Figure 1B). Based on this analysis, we designate these three clusters as matrix CAFs, chemokine CAFs, and contractile CAFs.

**Figure 1.**
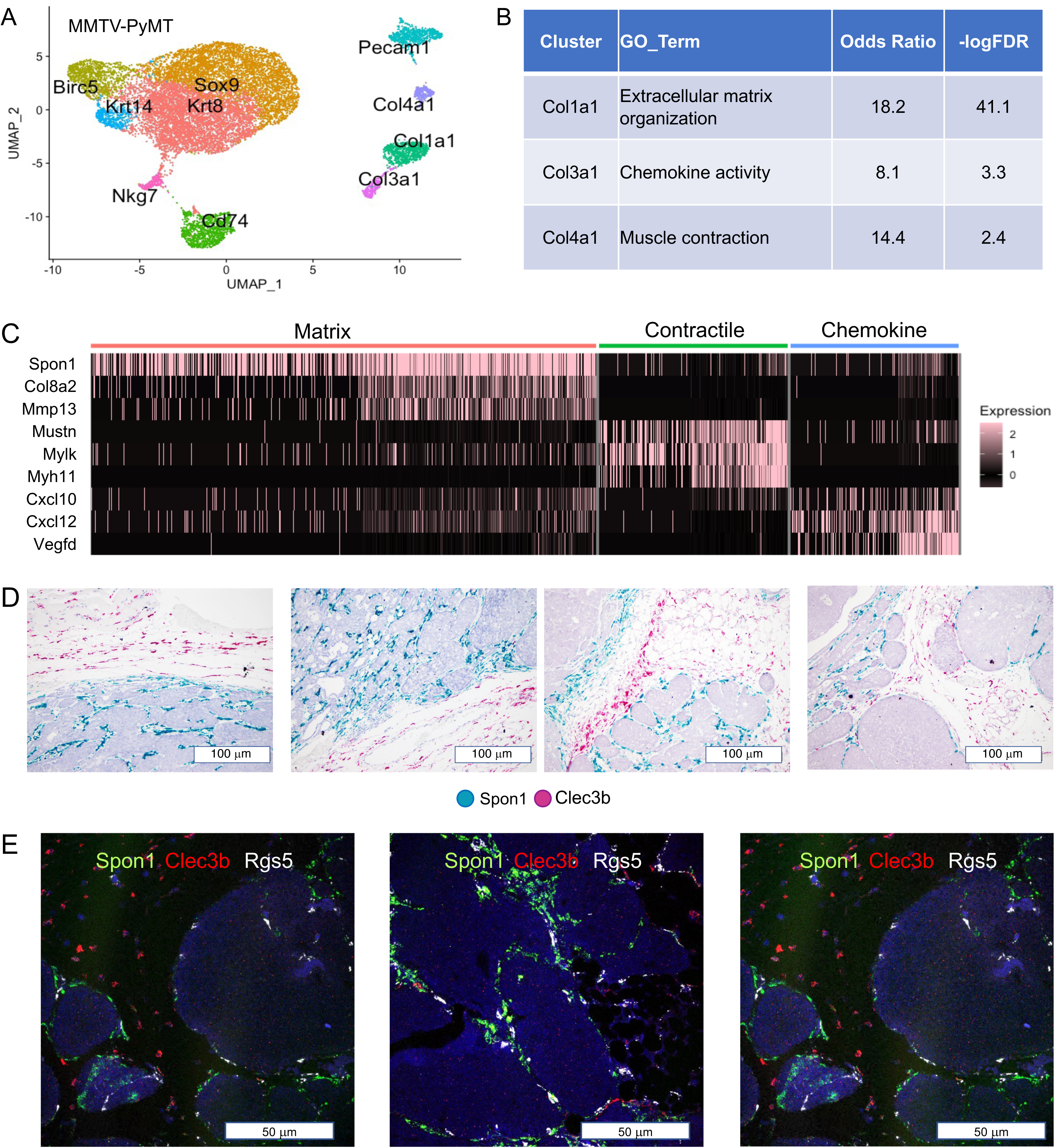
Single-cell RNA sequencing (scRNA-seq) characterization of CAFs in the MMTV-PyMT model. (A) Uniform Manifold Approximation and Projection (UMAP) visualization of all clusters identified in MMTV-PyMT mammary tumors. (B) Gene Ontology (GO) terms associated with the three distinct CAF subtypes. (C) Heatmap illustrating differentially expressed genes within each of the three CAF subtypes. (D) Images depicting distinct in situ hybridization (ISH) probes targeting Spon1 and Clec3b in hematoxylin-stained MMTV-PyMT tumors. (E) Fluorescent multiplexed ISH localization of *Spon1, Clec3b*, and *Rgs5* signals in MMTV-PyMT tumor tissue, with DAPI staining to visualize tumor cells.

Expression of seven differentially expressed genes associated with each of the three cluster’s associated GO term are displayed in a heatmap (Figure 1C). The genes associated with matrix CAFs encode matrix components or matrix remodeling proteins (*Spon1*, *Col8a2*, *Mmp13*) (Figure 1C). The genes associated with chemokine CAFs include *Cxcl10* and *Cxcl12*, along with *Vegfd*, which encodes a member of the VEGF-family that induces migration of lymphatic endothelial cells and peritumoral lymphangiogenesis ^11^ (Figure 1C). Contractile CAFs selectively express genes encoding components of the actomyosin cytoskeleton (*Myl11*, *Mylk*, *Mustn1*) (Figure 1C).

Our three CAF subtypes classification is in close agreement with three CAF subtypes revealed by scRNA-seq in melanoma described by Teichmann’s group (desmoplastic, immune, and contractile) ^12^ as well as the three CAF subtypes identified by scRNA-seq in MMTV-PyMT mammary tumors by Gallego-Ortega’s group (extracellular matrix, immune, and myofibroblast)^13^.

To determine the physical localization of the three different CAFs in the tumor microenvironment, we looked for genes which were expressed at high levels and strongly specific for one specific CAF subtype. The top markers for the three CAFs were *Spon1* (matrix CAFs), *Clec3b* (chemokine CAFs), and *Rgs5* (contractile CAFs) (Supplementary Figure 1A). Although *Rgs5* is sometimes considered as a definitive marker for pericytes and used to exclude cells from downstream analysis of CAFs ^6,14^, contractile CAFs expressed a multitude of fibroblast markers. Additionally, this class of fibroblastic cells has been previously designated as CAFs and shown to be localized away from the vasculature in both breast and melanoma tumors ^5,7,12^.

### Matrix CAFs are tumor-adjacent whereas chemokine CAFs are confined to the stroma

Having identified markers that are selectively expressed in the three CAF subtypes, our next step was to use these selective markers to determine their location within tumor tissue. We used RNA in situ hybridization (ISH) to visualize *Spon1* and *Clec3b* RNA molecules in hematoxylin-stained, formalin-fixed tissue samples of mammary tumors from MMTV-PyMT mice. The RNAscope probes to *Spon1* (matrix CAFs) and *Clec3b* (chemokine CAFs) were differentially labeled with a cyan dye and magenta dye, respectively. Shown in Figures 1D and S1B are twelve images taken from three different tumors, where matrix CAFs and chemokine CAFs display non-overlapping, discrete locations within the tumor. Matrix CAFs are located adjacent to the epithelial tumor nests, whereas chemokine CAFs are located within the surrounding stroma. In areas containing stromal adipocytes, *Clec3b* RNA staining is observed within some adipocytes, consistent with previous observations of its expression in murine adipose tissue ^15^. We note that due to their large size and other properties adipocytes are not robustly profiled by standard scRNA-seq methodology.

We quantified the localization of matrix CAFs and chemokine CAFs to areas containing tumor epithelial cells as opposed to stromal areas that were devoid of tumor cells. *Spon1* (matrix CAFs) staining comprised up to 6% of the total area containing tumor epithelial cells, and less than 1% of the total stromal area (p < 0.001); whereas *Clec3b* (chemokine CAFs) staining comprised up to 1.5% of the stromal area, but less than 0.2% of the tumor epithelial area (p < 0.001) (Figure S1C).

We detected corresponding physical separation of matrix CAFs and chemokine CAFs in MMTV-Erbb2 mammary tumors (Figure S1D). We also detected *Clec3b* and *Spon1* signals in normal mouse mammary tissue, with the *Spon1* signal coming from cells in between the two ducts and *Clec3b* signals being more distal (Figure S1E).

### Contractile CAFs are tumor-adjacent and intermingle with matrix CAFs

The first indication that contractile CAFs were intermingled with matrix CAFs was based on indirect evidence. RNAscope analysis of tumor tissue revealed the presence of CAFs intermingled with *Spon1*-expressing matrix CAFs that did not express the matrix CAF marker *Spon1* but clearly expressed *Acta2 or Pdgfrb*, genes that are highly expressed in contractile CAFs (Figure S1F). This intermingling was confirmed by multiplexed fluorescent RNA-ISH, using differentially labeled probes for all three CAF markers. As shown in Figure 1E, both *Rgs5* (contractile CAFs) and *Spon1* (matrix CAFs) expression was detected in between tumor cell nests, which appear dark blue from 4′,6-diamidino-2-phenylindole (DAPI) staining. There was more *Spon1* signal than *Rgs5*, consistent with the greater cell numbers of *Spon1*-expressing matrix CAFs relative to *Rgs5*-expressing contractile CAFs estimated by scRNA-seq (Figure 1E). Migratory CAFs were distally located away from the tumor nests as revealed by *in situ* fluorescence with *Clec3b* probe (Figure 1E).

### EMT transition in murine C3-SV40 T-ag breast tumors and analysis of CAFs

In parallel analyses using transgenic mouse models of breast cancer, C3(1)/SV40 T-antigen (C3-SV40-T) mice exhibited gene expression patterns resembling human basal breast cancer with mesenchymal features, in contrast to the luminal breast cancer features observed in MMTV-PyMT mice. Single-cell analysis of tumors from four C3-SV40-T mice revealed seven clusters, including immune cell clusters, stromal cell types and tumor cells. Fibroblasts constituted only 3% of total cells, significantly lower than in MMTV-PyMT tumors (Figure 2A). Two types of *Krt14*-expressing tumor epithelial cells were identified, one with high *Epcam* expression and another with high *Vim* expression. Cells belonging to the *Vim* cluster have undergone a complete epithelial-to-mesenchymal transition (EMT), as evidenced by a lack of expression of the epithelial marker *Cdh1*, concomitant expression of *Vim,* and expression of the EMT transcription factor *Twist1* (Figure 2B). RNA in situ hybridization (ISH) confirmed co-expression of SV40 T-antigen RNA with either *Cdh1* or *Vim* RNA transcripts at different stages of tumor development. At 14 weeks, a large majority of tumor cells co-expressed RNA for SV40 T-ag along with cadherin 1 RNA (Figure 2C). A few isolated tumor cells expressing vimentin were also observed (Figure 2D). By 28 weeks there had been a major transformation, such that the outer periphery of tumor nests were comprised of cells that co-expressed SV40 T-ag RNA along with *Cdh1* RNA (Figure 2E), whereas the inner cell mass of tumor nests were comprised of cells that SV40 T-ag RNA along with *Vim* RNA (Figure 2F).

**Figure 2.**
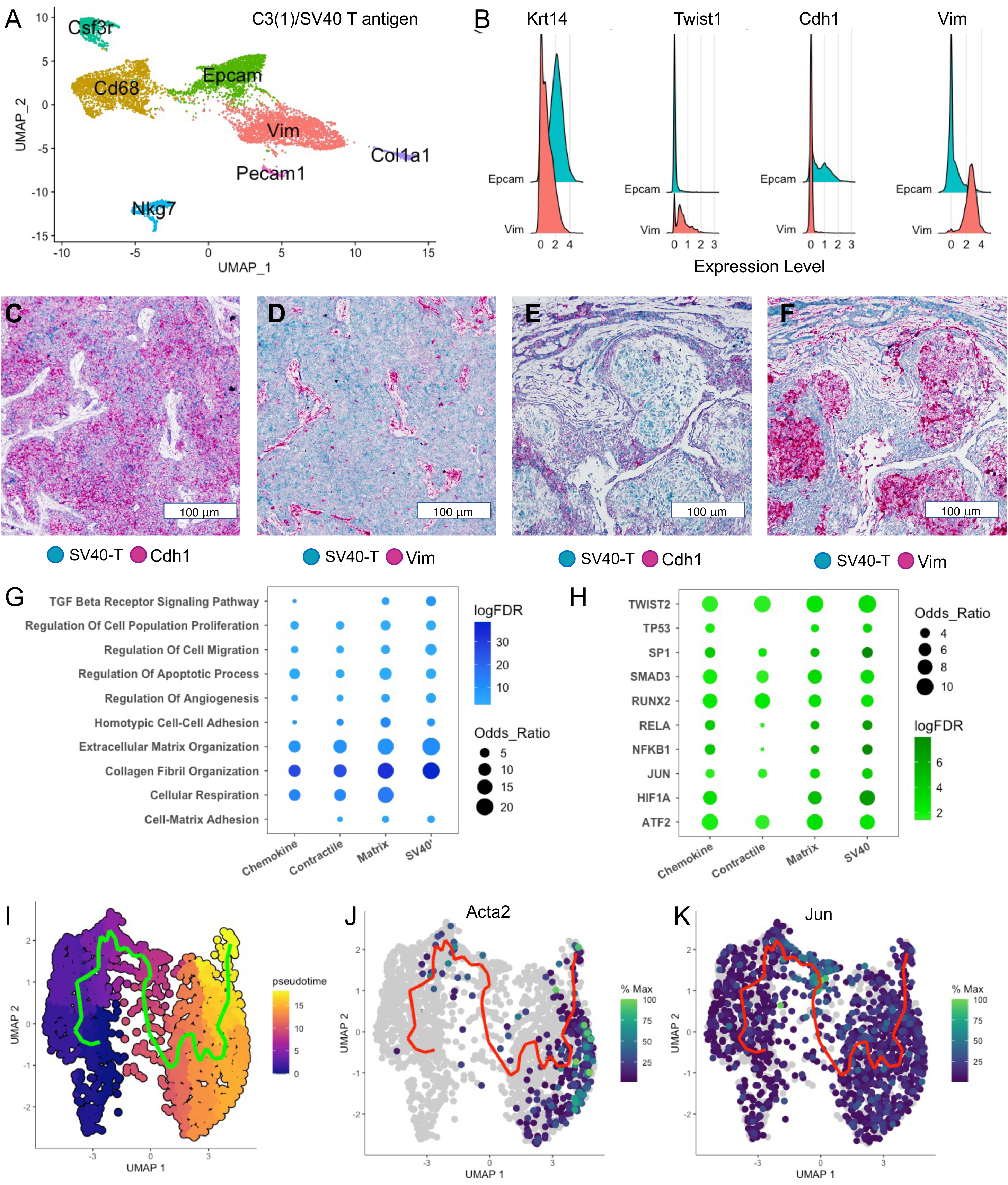
Comparative analysis between normal mammary stromal cells and CAFs in mouse breast cancer. (A) UMAP projection revealing all clusters identified in C3-SV40-Tag mammary tumors. (B) Plots comparing the expression of four genes in two distinct clusters of tumor cells: *Krt14* (myoepithelial marker), *Twist1* (EMT transcription factor), *Cdh1* (epithelial marker), and *Vim* (mesenchymal marker). (C, D) Images of 14-week-old C3-SV40-Tag mammary tumors, differentially labeled by ISH probes targeting SV40 and either *Cdh1* (C) or *Vim* (D). (E, F) Images of 28-week-old C3-SV40-Tag mammary tumors, differentially labeled by ISH probes targeting SV40 and either *Cdh1* (E) or *Vim* (F). (G) Dot plot displaying the extent and significance of GO term enrichment for biological processes in distinct CAFs. (H) Dot plot illustrating the extent and significance of enrichment for specific transcriptional target genes. (I) UMAP-based pseudotime trajectory, depicting the conversion of normal fibroblasts into matrix CAFs. (J) Expression of *Acta2* relative to the pseudotime trajectory. (K) Expression of *Jun* relative to the trajectory.

Although there was a single cluster of CAFs, further refinement generated two subclusters: one expressing high levels of *Clec3b*, the lead marker for migratory CAFs, and the other expressing higher levels of *Spon1*, the lead marker for matrix CAFs (Figure S2A). We observed spatial separation of these two subclusters in C3-SV40-T breast tumors, similar to MMTV-PyMT breast tumors (Figure S2B). The absence of *Rgs5* -expressing contracile CAFs suggests that some cancer-cells that have undergone EMT may fulfill functions carried out by contractile CAFs in MMTV-PyMT breast tumors.

### Comparing CAFs to their normal counterparts

Our study aimed to elucidate expression changes in cancer-associated fibroblasts (CAFs) during cancer progression. While the exact origin of most CAFs remains unclear, existing evidence suggests derivation from tissue-resident cells^1,2,16^. To compare CAFs with normal breast tissue-resident cells, we performed scRNA-seq analysis on mammary glands from mice genetically matched to MMTV-PyMT and C3-SV40-Tag mammary tumors. Differential gene expression analysis revealed seventeen clusters, including epithelial compartments (luminal, myoepithelial, and developmental), six immune cell clusters, three endothelial cell clusters, pericytes (*Rgs5*), and three *Col1a1+* fibroblastic clusters expressing selective markers (*Spon1*, *Clec3b*, and *C3*). The *Rgs5* cluster exhibited canonical pericyte markers and additional pericyte markers obtained from the Enrichr database^17^.

To characterize the three normal mammary tissue fibroblastic clusters, we compared their top one-hundred differentially expressed genes (ranked by FDR) to pan-tissue fibroblast lineages identified in a comprehensive scRNA-seq study across twenty normal tissues. These lineages included *Pi16+* and *Col15a1+* found ubiquitously and *Ccl19+* found in spleen and lymph nodes^18^. Remarkably, over 80% of the top differentially expressed genes in our *Clec3b+* cluster matched those of the *Pi16+* lineage, known as the most stem-like fibroblast lineage (Figure S2E). Similarly, the other two mammary fibroblast clusters showed over 80% identical matches with markers of the three fibroblast lineages: *Spon1+* fibroblasts matched *Pi16+* and *Col15a1+* lineages equally, while *C3+* fibroblasts predominantly matched the *Ccl19+* lineage (Figure S2E).

Using Seurat, we constructed an integrated dataset of tissue-resident fibroblasts and pericytes from normal mammary glands, along with CAFs from MMTV-PyMT and C3-promoter driven SV40 T-antigen mammary tumors. There were four fibroblastic clusters in the integrated dataset, including a cluster labelled “Lymph Node” that was comprised solely of normal mammary cells (Figure S2F). MMTV-PyMT tumors contained a greater percentage of CAFs (9%) than C3-SV40 T-antigen tumors (3% CAFs). Additionally, MMTV-PyMT tumors had less chemokine fibroblasts and more contractile fibroblasts compared to normal mammary tissue, similar to the findings of Teichmann’s group in melanoma^12^ (Figure S2F).

We were interested to determine the degree to which canonical CAF markers were expressed in the different CAF subtypes, and how this expression differed from the expression In their corresponding normal cell type. For matrix CAFs, the expression of all five markers (*Pdgfra, Pdgfrb, Fap, Acta2, S100a4*) was increased in PyMT tumors (Figure S2H). For chemokine CAFs, *S100a4* was the CAF marker with the largest increase, and *Pdgfra* and *Pdgfrb* expression increased slightly (Figure S2H). Finally, *Pdgfrb* was the most differentially expressed marker in contractile CAFs although *S100a4* and *Acta2* showed more modest increases (Figure S2H).

### Gene sets overrepresented in the transition from normal tissue resident cells to CAFs

To gain a genome-wide perspective on the gene expression changes accompanying CAF cancer progression, we performed gene set enrichment analysis using the differentially (over)expressed genes in MMTV-PyMT and SV40-Tag CAFs along with GO term enrichment analysis for biological processes (http://geneontology.org). Most of the enriched biological processes have been previously shown to play a role in CAF development, including collagen fibril and extracellular matrix organization, and regulation of angiogenesis, cell-matrix adhesion, migration, cell proliferation, and TGF beta signaling (Figure 2G). To the best of our knowledge, regulation of apoptosis, cell-cell adhesion, and cellular respiration have not been previously linked to induction of CAFs.

To gain insight into what might be driving CAF development, we performed enrichment analysis of known gene targets for transcriptional regulators^19^. Several of the transcriptional regulators identified (*Twist2, Runx2, Smad3, Nfkb1, Rela, Hif1a, Tp53*) are known to be involved in CAF development ^20^. Two others (*Jun, Atf2*) encode subunits of the AP-1 transcriptional factor, which together with *Sp1* can play a key role in reprogramming ^21^.

### Trajectory analysis of the conversion of normal matrix fibroblasts into matrix CAFs

We next performed trajectory analysis with Monocle3 to explore whether there were discrete transcriptomic transitions during the conversion of normal fibroblasts into CAFs. We first focused on matrix CAFs, which by far showed the biggest alterations in gene expression during cancer progression (> five-fold more significantly altered genes than either migratory or contractile CAFs). We found a single pseudotime trajectory connecting two groups of cells with clearly different pseudotime values (Figure 2I). The expression of the CAF marker *Acta2* is closely aligned with advanced pseudotime values (Figure 2J). Interestingly, *Jun* expression is highest in the bridge of cells connecting the two large groups (Figure 2K).

We corroborated this transitory transcriptional state using an alternate trajectory inference method (scFates)^22^. Similar to our results using Monocle, a UMAP projection of the matrix fibroblasts showed two separate populations that were connected by the pseudotime trajectory (Figure 3A). While the expression of *Col1a1* increased in concert with pseudotime, the expression of Jun reached its zenith in the middle of the trajectory (Figure 3A).

**Figure 3.**
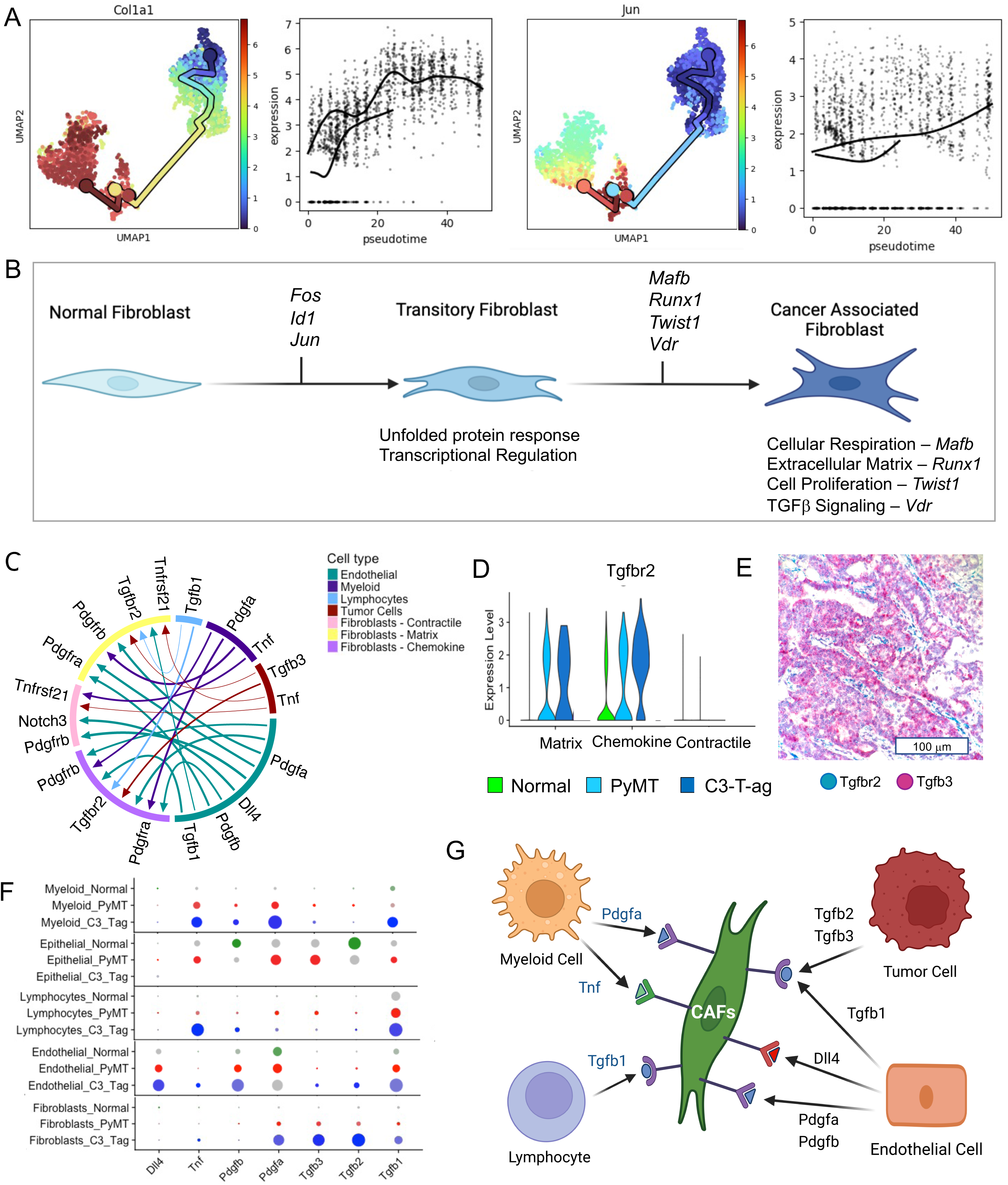
Analysis of transcriptional changes and tumor microenvironment induction in murine CAFs. (A) Expression of *Acta2* or *Jun* relative to scFates pseudotime trajectory depicting the conversion of normal fibroblasts into matrix CAFs. (B) Model depicting the transcriptional alterations during CAF development. (C) Circle plot illustrating the ligand-receptor interactions between CAFs and other cells within the tumor microenvironment. (D) Violin plots showing the expression of *Tgfrb2* in CAFs and normal fibroblasts. (E) Representative image of differentially labeled ISH probes targeting *Tgfbr2* and *Tgfb3* in hematoxylin-stained MMTV-PyMT tumors. (F) Dot plot displaying the expression of indicated ligands in various cell types within the tumor microenvironment. (G) Schematic representation of how different cells within the MMTV-PyMT tumor microenvironment utilize TGFβ, PDGF, Notch, and TNF ligands to mediate the activation of CAFs.

A useful feature of Monocle3 is the ability to find modules of co-regulated genes that share similar pseudotime values^8^. Of particular interest was the module (#23) that contained *Jun* and 101 other genes, 23 of which encode transcription factors including several components of the AP-1 transcriptional complex and 10 protein folding chaperones (Table S1). The average expression of the genes that comprise this module show a highly similar trajectory to *Jun* itself, with the highest expression along the bridge that connects the two large groups (Figure S3A).

We next examined modules that showed strong expression in the right major group containing CAFs. We reasoned that some of the biological processes associated with CAF formation might be caused by distinct transcriptional programs. In support of this hypothesis, we identified modules with similar expression patterns (Figure S3B and S3C) but with contrasting gene enrichments: one exhibited a significant enrichment of mitochondrial respiratory genes while lacking enrichment for extracellular matrix genes, whereas the other module displayed the opposite pattern (Table S1). Based on these findings, a model for how normal matrix fibroblasts are converted into matrix CAFs is shown in Figure 3B. A transitory state is marked by the expression of a large number of transcription factors presumably involved in the initial stages of reprogramming, including *Jun, Fos*, and *Id3*, followed by expression of transcription factors that induce distinct CAF phenotypes, such as *Mafb* associated with expression of mitochondrial respiratory genes, *Runx1* associated with extracellular matrix, *Twist1* associated with cell proliferation, and *Vdr* associated with TGFβ signaling (Figure 3B).

While we were successful in finding a single trajectory closely aligned with the conversion of normal matrix fibroblasts into matrix CAFs, we were unable to do so with either migratory or contractile CAFs.

### Reciprocal upregulation of pro-CAF ligands in the murine mammary tumor microenvironment along with upregulation of corresponding receptors in CAFs

To gain insight into how the tumor microenvironment induces the conversion of normal fibroblasts into CAFs, we utilized LIANA which aggregates the results from combination of ligand-receptor analysis methods and resources^23^. Restricting our attention to signal transduction receptors present on CAFs and their associated ligands, we filtered this set for cases where receptors and ligands were both overexpressed in PyMT tumors relative to normal cells. This yielded six ligand-receptor pairs, and the microenvironment cells producing the ligand and corresponding CAF receptors are shown in Figure 3C.

We found that the receptor gene *Tgfbr2* was upregulated in both matrix and chemokine CAFs (Figure 3D), and that a corresponding ligand gene *Tgfb3* was upregulated in tumor cells (Figure 3F). To corroborate this reciprocal signaling system, we used RNA in situ hybridization (ISH) to visualize *Tgfbr2* and *Tgfb3* RNA molecules in mammary tumors from MMTV-PyMT mice. The RNAscope probes to *Tgfbr2* and *Tgfb3* were differentially labeled with a cyan dye and magenta dye, respectively. Shown in Figure 3E is a representative image of the magenta *Tgfb3* probe brightly staining the tumor epithelial cells, whereas the cyan *Tgfbr2* probe is staining the fibroblasts lying in between tumor cell nests. By applying NicheNet, a computational method that predicts ligand–target links between interacting cells by testing whether gene sets associated with the response to a given ligand are induced in a target cell expressing the corresponding receptor ^10^, we found that that the many of the known transcriptional targets of TGF-beta receptor ligands were upregulated in matrix CAFs, establishing this reciprocal signaling system (Figure S3D).

The other reciprocal ligand-receptor interactions we observed included the ligand gene *Dll4* activated in endothelial cells with corresponding activation of its receptor gene *Notch3* specifically in contractile CAFs, the ligand gene *Tnf* activated in tumor and myeloid cells coupled with activation of the receptor gene *Tnfrsf21* in matrix and contractile CAFs, and activation of *Pdgfa* and *Pdgfb* in endothelial and myeloid cells and *Tgfb1b* in lymphocytes (Figures 3F and S3E). Based on these findings, a model for how secretion of these four classes of secreted ligands in the MMTV-PyMT mammary tumor microenvironment stimulates the development of CAFs through corresponding receptors is shown in Figure 3I.

### Human breast cancers exhibit three CAF subtypes analogous to murine CAF subtypes

To compare our results from mouse models to human breast cancer, we performed scRNA-seq analysis of three human breast cancer samples and integrated these using Seurat along with three normal human mammary datasets. We found seven epithelial cell clusters, two immune cell clusters, and one endothelial cell cluster. The most striking difference observed difference between the normal and tumor samples was the near complete disappearance of *KRT14* and *KRT17* myoepithelial cells in tumors (Figure 4A,4B). We observed three collagen expressing clusters labeled *COL1A1*, *DCN*, and *RGS5* (Figure 4A, ,4B). The results of GO term enrichment analysis of genes selectively expressed in the each of the CAF clusters was highly similar to results obtained with murine CAFs: the three groups were enriched for extracellular matrix organization, chemokine activity, and muscle contraction, respectively (Figure 4C).

**Figure 4.**
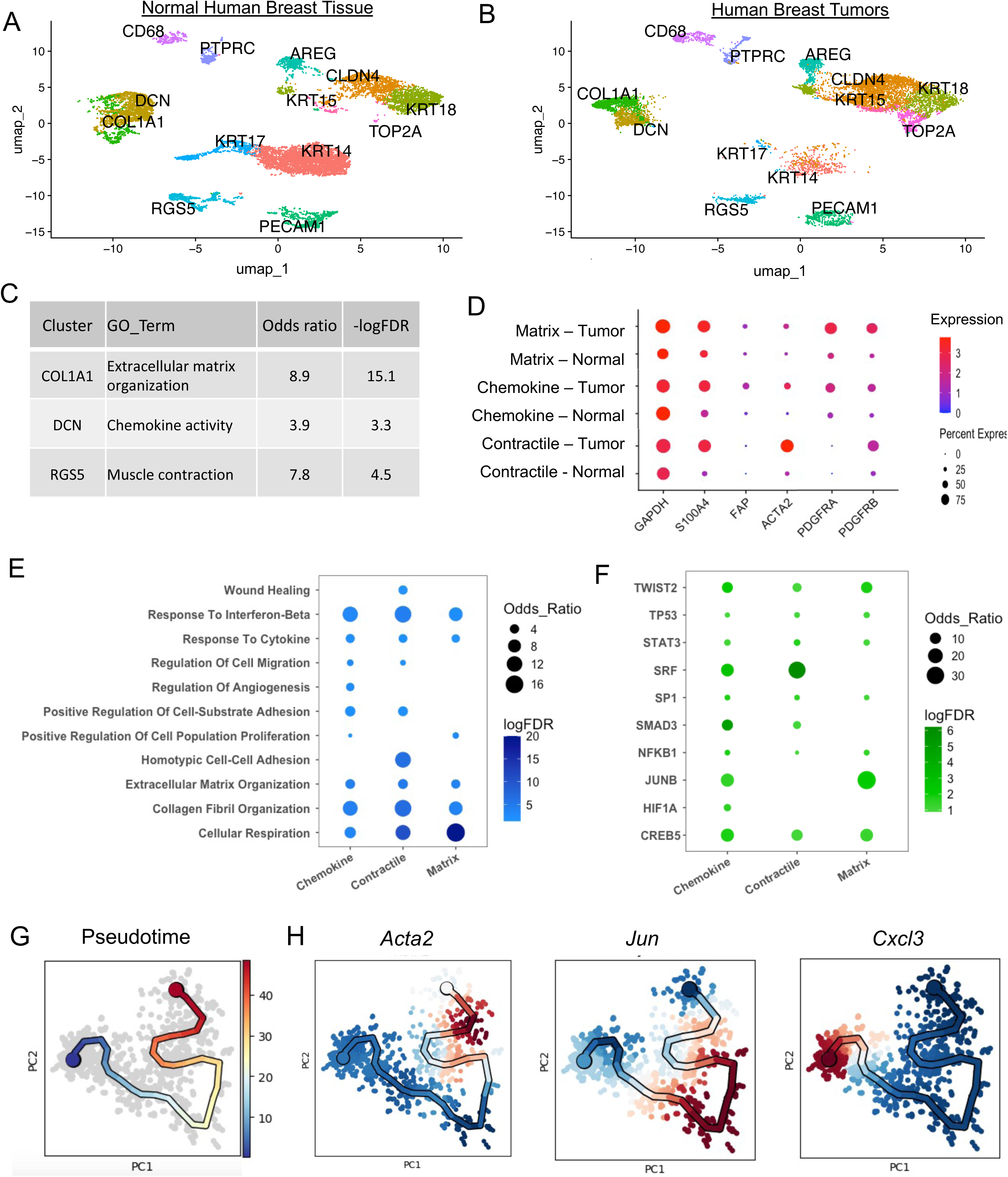
Comparative analysis between normal mammary stromal cells and human breast CAFs. (A) UMAP projection of transcriptional clusters identified from scRNA-seq analysis of human normal mammary tissue. (B) UMAP projection of transcriptional clusters identified from scRNA-seq analysis of human breast cancers. (C) GO terms associated with the three distinct CAF subtypes. (D) Dot plot showing the differential expression of canonical CAF markers. (E) Dot plot illustrating the extent and significance of GO term enrichment for biological processes in different human CAFs. (F) Dot plot illustrating the extent and significance of enrichment for specific transcriptional target genes. (G) UMAP-based scFates pseudotime trajectory depicting the conversion of normal cells into contractile CAFs. (H) Expression of *Acta2, Jun*, and *Cxcl3* relative to the scFates pseudotime trajectory.

Following our approach with murine CAFs, we investigated the expression levels of canonical CAF markers across different CAF subtypes and compared them with normal cells. In matrix CAFs, we observed upregulation of three markers (*PDGFRA, PDGFRB, S100A4*) in tumor cells, accompanied by a modest increase in *FAP* and *ACTA2* expression (Figure 4D). Similarly, chemokine CAFs exhibited comparable trends, albeit with higher induction levels of *FAP* and *ACTA2*, and less induction of *PDGFRA* and *PDGFRB* (Figure 4D). Notably, contractile CAFs displayed robust upregulation of *S100A4*, *ACTA2* and *PDGFRB* (Figure 4D).

We conducted gene set enrichment analysis using differentially expressed genes in human CAFs, along with GO term enrichment analysis for biological processes, mirroring our methodology with murine CAFs. Many biological processes enriched in murine CAFs, such as cellular respiration, collagen fibril and extracellular matrix organization, and regulation of angiogenesis, cell-substrate adhesion, migration, and cell proliferation, were similarly observed in human CAFs (Figure 4E). Unique to human CAFs were responses to cytokines and interferon-beta (Figure 4E). Enrichment analysis of known gene targets for transcriptional regulators was conducted. Among the identified transcriptional regulators, several we previously found in murine CAF development (*TWIST2, SMAD3, NFKB1, HIF1A, TP53*). Additionally, human CAFs exhibited unique involvement of *STAT3* and *CREB5* (Figure 4F).

### Trajectory analysis of the conversion of normal pericytes into contractile matrix CAFs

We performed trajectory analysis using the tumor-normal integrated data and were able to detect an intermediary state characterized by Jun and Fos expression in contractile CAFs, but failed to detect a transitional state in the other two other CAFs. Possible reasons for this discrepancy are outlined in the discussion. The pseudotime trajectory for contractile CAFs was computed using scFates with *Cxcl3* as the root or starting point as it is selectively expressed in normal pericytes (Figure 3G). By plotting the expression levels of *Acta2*, *Jun*, and *Cxcl3* along this trajectory it is apparent that expression of *Jun* is greatest in the middle of the pseudotime trajectory, whereas *Cxcl3* expression is greatest at the beginning point and *Acta2* is greatest at the end point (Figure 3H).

By first filtering for genes that were significantly associated with pseudotime and then performing cluster analysis, we found gene expression modules that contained genes with a similar trajectory relationships, two of which are displayed in Figure 5A. On the left are genes with maximal expression towards the pseudotime endpoint, including the smooth muscle genes *Myl9* and *Myh11*. On the right are genes with highest expression in the middle of the trajectory, including *Junb, Fos* and several protein chaperone genes (Figure 5A). These results show evolutionary conservation of a mechanism underlying CAF development. However, we did not detect a transitory state for either matrix CAFs or chemokine CAFs. In matrix CAFs, Jun expression has the same pseudotime association at the endpoint as does *COL1A1* or *PDGFRB* (Figure 5B).

**Figure 5.**
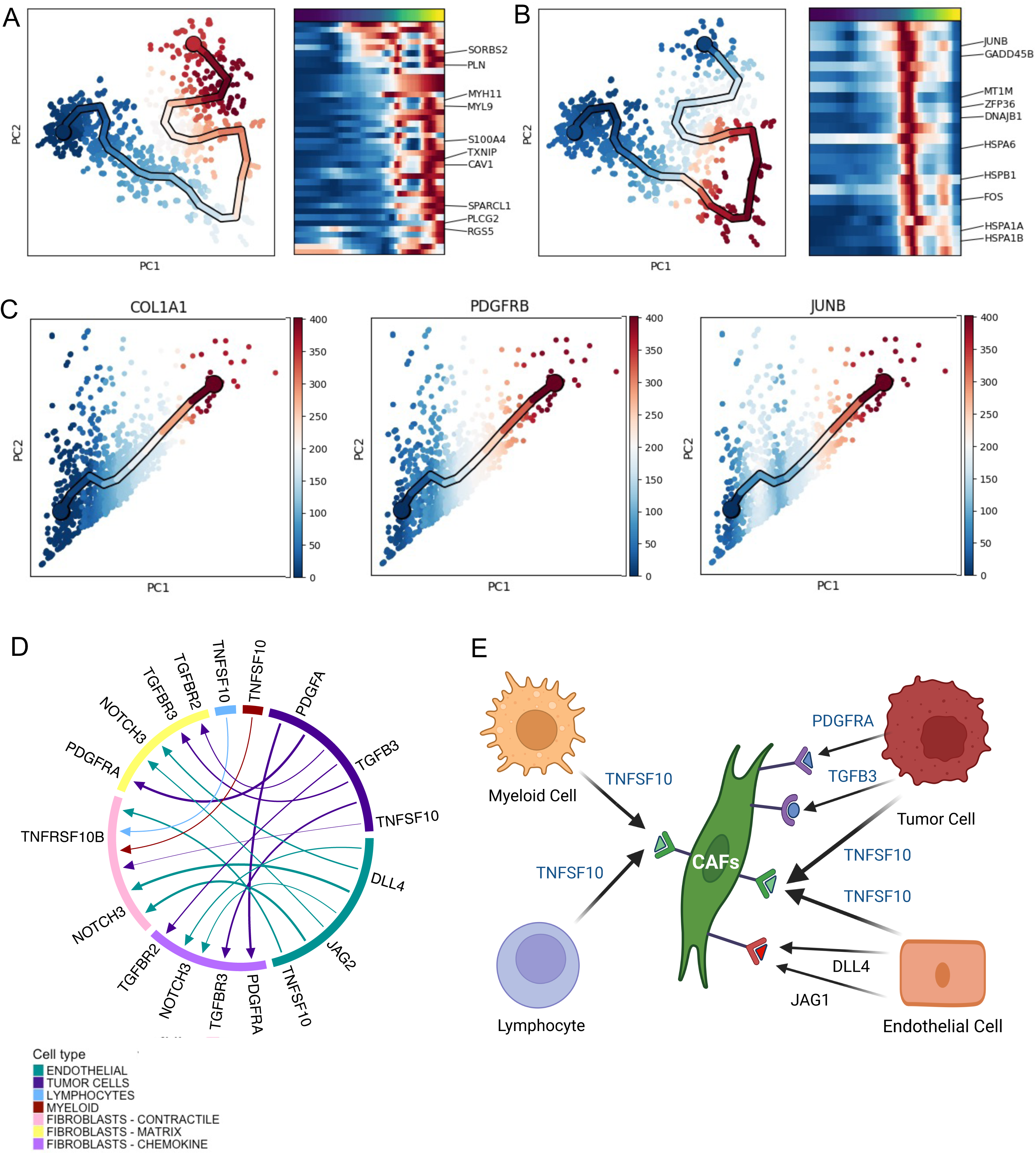
Analysis of transcriptional changes and tumor microenvironment induction in humanCAFs. (A) Aggregated expression genes belonging to a module linked to the endpoint of the pseudotime trajectory of the conversion of normal fibroblasts into contractile CAFs, accompanied by a heatmap displaying associated genes. (B) Expression of a different gene module linked to the transitory state of the conversion of normal fibroblasts into contractile CAFs, accompanied by a heatmap displaying associated genes. (C) Expression of *Col1a1, Pdgfrb*, and *Jun* relative to the trajectory of the conversion of normal fibroblasts into matrix CAFs. (D) Circle plot illustrating the ligand-receptor interactions between CAFs and other cells within the tumor microenvironment. (E) Schematic representation of how different cells within the human breast cancer tumor microenvironment utilize TGFβ, PDGF, Notch, and TNF ligands to mediate the activation of CAFs.

### Reciprocal upregulation of pro-CAF ligands in the human mammary tumor microenvironment along with upregulation of corresponding receptors in CAFs

To test if the tumor microenvironment in human breast cancer induces the conversion of normal fibroblasts into CAFs in a manner similar to mouse model breast cancer, we utilized LIANA to find ligand-receptor pairs and restricted our attention to signal transduction receptors present on CAFs and their associated ligands, such that the receptors and ligands were both overexpressed in cancerous relative to normal mammary tissue. This yielded five ligand-receptor pairs, three of which were identical to ligand-receptor pairs observed in the MMTV-PyMT model and two of which were analogous. The microenvironment cells producing the ligand and corresponding CAF receptors are shown in Figure 5F. Ligand-receptor expression results are shown in Figure S4A-C. As we found in previously in the MMTV-PyMT mouse model, all cells within the tumor microenvironment contributed to CAF education. A model for these results in shown in Figure 5G.

## Discussion

The two primary goals of our study were to uncover new insights into cancer-associated fibroblast (CAF) development and to assess the validity of established features in this process. We identified previously undescribed aspects of CAF development, including a significant enrichment of cellular respiratory gene expression in both human and mouse CAFs. While increased cellular respiration primarily generates ATP, our findings suggest that heightened flux through the mitochondrial Krebs cycle may also provide metabolic intermediates crucial for cancer cell growth, resembling the growth-promoting mechanism proposed for increased glycolysis in CAFs^2^.

Furthermore, we observed a potential transitory transcriptional state during CAF development, characterized by the expression of specific transcription factors and protein chaperones. This transient state shares similarities with observations in other cellular differentiation processes, including stage-specific expression of *Jun* and *Fos* genes during epidermal differentiation and the transient accumulation of the chaperone Hsp27 during differentiation of leukemia cells. suggesting a dynamic regulatory mechanism at play. Although there are other possible explanations, it may be that for the cases where we did not observe a transitory state in the development of CAFs that full differentiation into mature CAFs had already occurred, which would be consistent with our analysis of human matrix CAFs (Figure 5C).

Our study also confirmed many previously described aspects of CAF development, supporting the selective expression of canonical CAF markers due to tumorigenesis rather than originating cell properties. Additionally, we found concordance with established biological processes linked to CAF formation, such as extracellular matrix organization, cellular migration, wound healing and the TGFβ signaling pathway. Furthermore, our findings strengthen previous research implicating PDGF, TGF-β, Notch, and TNF secreted ligands in the formation of CAFs. Moreover, we offered corroborating evidence that multiple, if not all, cell types within the tumor microenvironment contribute to CAF formation.

In our investigation of breast cancer CAFs, we identified three distinct subtypes (matrix, chemokine, and contractile) in both human and murine samples. These findings align closely with previous reports, including one that also featured the MMTV-PyMT model and identified three CAF subtypes (desmoplastic, inflammatory, and contractile) containing marker genes that overlapped with our subtypes^13^. Our results with are also in agreement with analysis of human breast cancer CAFs by Swarbrick’s group, who found four groups: myCAFs , corresponding to matrix CAFs; iCAFs, corresponding to chemokine CAFs, and two perivascular groups corresponding to contractile CAFs. Importantly, they observed that their iCAFs were located distal to tumor cells, consistent with our finding that chemokine CAFs are tumor-distal ^7^.

Overall, our study enhances our understanding of CAF development by uncovering novel mechanisms and validating established features including the classification of breast cancer CAF subtypes. These findings have implications for advancing our understanding of the role of CAFs in cancer progression and may serve as a foundation for future basic research aimed at unraveling the complexities of the tumor microenvironment.

## Methods

### Animals

Animal experiments were carried out at the Division of Laboratory Animal Resources at Stony Brook University in accordance with Institutional Animal Care and Use Committee-approved procedures. Wild-type (FVB) and MMTV-PyMT (FVB) mice were obtained from Jackson Laboratory. Additional MMTV-PyMT (FVB) and the C3(1)/Tag (FVB) mice were obtained from Dr. Mikala Egeblad (Cold Spring Harbor Laboratory). Four 12-week-old MMTV-PyMT/FVB mice, two corresponding wild-type FVB mice, and four C3(1)/Tag (FVB) mice were used to prepare the mammary tumor single-cell preparations for scRNA-seq and histological slides. In addition, tissue sections from mammary glands of a 52-week Erbb2 transgenic mouse obtained from Dr. Natasha Marchenko were used for histological studies.

### Tumor sample dissociation into a single-cell suspension

The mammary tumors were harvested from the mice’s left and right mammary glands (four and five). The tissue was finely minced and placed in 15 ml conical tubes with a dissociation solution composed of Collagenase/Hyaluronidase (Stem Cell Technologies) enzymes, 5% fetal bovine serum (FBS) (Gemini Bioproducts), and 1 mg/mL DNase 1 (Stem Cell Technologies). The samples were digested for 1 hour at 37°C with constant low agitation using a thermomixer (Eppendorf). The digested cells were pelleted and washed twice with RPMI media in the presence of 5% FBS. The cell suspensions were filtered using a 70 µm cell strainer (DB) to collect single-cell suspensions and single-cell digested tissue. The cells were resuspended and washed twice in 3 ml Red Blood Cell Lysis Buffer (Roche) to remove any visible blood cells. The final cell suspensions were pelleted by centrifuging at 300Xg for 5 minutes and resuspended in 1 ml RPMI media with 5% FBS and restrained again with a 70 µm strainer to a final concentration of 10,000 cells/ml. The cell viability was examined using trypan blue exclusion (Invitrogen).

### scRNA-seq and sequencing library construction

Approximately 10,000 single cells resulting from the single-cell suspensions of each tumor were loaded into the 10x Genomics microfluidics device along with 10X Genomics gel beads (kit v2 PN-120237) containing barcoded oligonucleotides, reverse transcription (RT) reagents, and oil, resulting in gel beads in emulsion. The scRNA-seq library preparation followed the manufacturer’s protocol (10x Genomics) using the Chromium Single Cell 3-Library. These libraries were paired end sequenced using the Illumina HiSeq 4000 sequencing system.

### Computational analysis

We used Cell Ranger to convert single-cell sequencing files into a data structure suitable for Seurat. Seurat was used for quality control and preprocessing, dimensionality reduction, cell clustering and annotation, integration of multiple datasets, and differential expression analysis. We performed trajectory analysis using Monocle3 and scFates. Cell-cell communication was analyzed using Liana and Nichenet. Gene set enrichment was carried out with the Gene Ontology Resource and Enrichr.

### Tissue preparation of histological sectioning, fixation and staining

The tissue sections preserved for histology purposes fixed in formalin, were switched to 70% ethanol within 24 hours and were sent to the Stony Brook Histology Core for paraffin embedding. Tissue sections were cut to 5 μm thickness and were assessed by RNAScope. Images were captured at 20X and 40X magnification.

### RNAScope mRNA In Situ Hybridization Assay

RNAscope® technology was used to perform the assay to spatially resolve our scRNA-seq data. The 2.5 HD Duplex Reagent Kit (CN: 322430, Advanced Cell Diagnostics) and RNAscope® Fluorescent Multiplex Reagent Kit (CN: 320850, Advanced Cell Diagnostics) were used to perform the chromogenic and fluorescent ISH assays respectively. The mRNA expression of our gene markers was detected using the following murine probes: RNAscope® Probe – Mm-Col1a1 (CN: 319371), RNAscope® Probe – Mm-Clec3b C2 (CN: 300031-C2), RNAscope® Probe – Mm-Spon1 (CN: 300031), RNAscope® Probe – Mm-Pdgfrβ (CN: 411381), RNAscope® Probe – Mm-Acta2 (CN: 319531), RNAscope® Probe – Mm-Fap (CN: 423881), RNAscope® Probe – Mm-Ccl7 (CN: 446821), RNAscope® Probe – Mm-Krt14-C2 (CN: 422521-C2), RNAscope® Probe – Mm-Cxcl12 (CN: 422711), RNAscope® Probe – Mm-Cxcl14 (CN: 459741), RNAscope® Probe – Mm-Cxcr4-C2 (CN: 425901-C2), RNAscope® Probe – Mm-Ccr1-C2 (CN: 402721-C2), RNAscope® Probe – Mm-Rgs5-C3 (CN: 430181-C3). All tissues treated for chromogenic ISH were counterstained using Mayer’s Hematoxylin.

## Data Deposition and Access

Single-cell sequencing data generated in the course of this study will be available in NCBI GEO datasets upon publication. To review GEO accession GSE199515, go to https://www.ncbi.nlm.nih.gov/geo/query/acc.cgi?acc=GSE199515 and enter the token mravkaqurxejtin into the box.

## Acknowledgements

This project was supported by funding from the NCI (R01 CA217206 to S. Powers) and the National Human Genome Research Institute (R21 HG009255 to S. Powers). The authors thank members of the Powers, Egeblad and Moll labs for discussions. The authors also thank the staff in the Research Histology Core and the Division of Laboratory Animal Resources (DLAR) in the Stony Brook School of Medicine for their technical support, as well as the staff in the Tissue Analytics, Biostatistics & Data Science Shared Resources of the Stony Brook Cancer Center.

## Author Contributions

APD performed the mouse experiments, sample/library preparation, histology staining and RNAscope analysis with assistance from AN, ME and UMM. ADP, BN and SP performed computational analysis of the data and interpreted the results, with contributions from JL and JP. SP and ME conceived and directed the study. NM provided critical tissue samples, SP wrote the first draft of the manuscript using text and figures prepared by APD. APD revised the first draft. All authors read and accepted the manuscript.

## Notes

### Competing Interest Statement

The authors have declared no competing interest.

### Summary of Updates

Based on reviewer comments, this manuscript has been extensively revised

## References

1. Arina, A., Idel, C., Hyjek, E.M., Alegre, M.L., Wang, Y., Bindokas, V.P., Weichselbaum, R.R., and Schreiber, H. (2016). Tumor-associated fibroblasts predominantly come from local and not circulating precursors. Proc Natl Acad Sci U S A 113, 7551–7556. 10.1073/pnas.1600363113.

2. Sahai, E., Astsaturov, I., Cukierman, E., DeNardo, D.G., Egeblad, M., Evans, R.M., Fearon, D., Greten, F.R., Hingorani, S.R., Hunter, T., et al. (2020). A framework for advancing our understanding of cancer-associated fibroblasts. Nat Rev Cancer 20, 174–186. 10.1038/s41568-019-0238-1.

3. Kalluri, R. (2016). The biology and function of fibroblasts in cancer. Nat Rev Cancer 16, 582–598. 10.1038/nrc.2016.73.

4. Beacham, D.A., and Cukierman, E. (2005). Stromagenesis: the changing face of fibroblastic microenvironments during tumor progression. Semin Cancer Biol 15, 329–341. 10.1016/j.semcancer.2005.05.003.

5. Bartoschek, M., Oskolkov, N., Bocci, M., Lovrot, J., Larsson, C., Sommarin, M., Madsen, C.D., Lindgren, D., Pekar, G., Karlsson, G., et al. (2018). Spatially and functionally distinct subclasses of breast cancer-associated fibroblasts revealed by single cell RNA sequencing. Nat Commun 9, 5150. 10.1038/s41467-018-07582-3.

6. Friedman, G., Levi-Galibov, O., David, E., Bornstein, C., Giladi, A., Dadiani, M., Mayo, A., Halperin, C., Pevsner-Fischer, M., Lavon, H., et al. (2020). Cancer-associated fibroblast compositions change with breast cancer progression linking the ratio of S100A4(+) and PDPN(+) CAFs to clinical outcome. Nat Cancer 1, 692–708. 10.1038/s43018-020-0082-y.

7. Wu, S.Z., Roden, D.L., Wang, C., Holliday, H., Harvey, K., Cazet, A.S., Murphy, K.J., Pereira, B., Al-Eryani, G., Bartonicek, N., et al. (2020). Stromal cell diversity associated with immune evasion in human triple-negative breast cancer. EMBO J 39, e104063. 10.15252/embj.2019104063.

8. Cao, J., Spielmann, M., Qiu, X., Huang, X., Ibrahim, D.M., Hill, A.J., Zhang, F., Mundlos, S., Christiansen, L., Steemers, F.J., et al. (2019). The single-cell transcriptional landscape of mammalian organogenesis. Nature 566, 496–502. 10.1038/s41586-019-0969-x.

9. Trapnell, C., Cacchiarelli, D., Grimsby, J., Pokharel, P., Li, S., Morse, M., Lennon, N.J., Livak, K.J., Mikkelsen, T.S., and Rinn, J.L. (2014). The dynamics and regulators of cell fate decisions are revealed by pseudotemporal ordering of single cells. Nat Biotechnol 32, 381–386. 10.1038/nbt.2859.

10. Browaeys, R., Saelens, W., and Saeys, Y. (2020). NicheNet: modeling intercellular communication by linking ligands to target genes. Nat Methods 17, 159–162. 10.1038/s41592-019-0667-5.

11. Tammela, T., and Alitalo, K. (2010). Lymphangiogenesis: Molecular mechanisms and future promise. Cell 140, 460–476. 10.1016/j.cell.2010.01.045.

12. Davidson, S., Efremova, M., Riedel, A., Mahata, B., Pramanik, J., Huuhtanen, J., Kar, G., Vento-Tormo, R., Hagai, T., Chen, X., et al. (2020). Single-Cell RNA Sequencing Reveals a Dynamic Stromal Niche That Supports Tumor Growth. Cell Rep 31, 107628. 10.1016/j.celrep.2020.107628.

13. Valdes-Mora, F., Salomon, R., Gloss, B.S., Law, A.M.K., Venhuizen, J., Castillo, L., Murphy, K.J., Magenau, A., Papanicolaou, M., Rodriguez de la Fuente, L., et al. (2021). Single-cell transcriptomics reveals involution mimicry during the specification of the basal breast cancer subtype. Cell Rep 35, 108945. 10.1016/j.celrep.2021.108945.

14. Elyada, E., Bolisetty, M., Laise, P., Flynn, W.F., Courtois, E.T., Burkhart, R.A., Teinor, J.A., Belleau, P., Biffi, G., Lucito, M.S., et al. (2019). Cross-Species Single-Cell Analysis of Pancreatic Ductal Adenocarcinoma Reveals Antigen-Presenting Cancer-Associated Fibroblasts. Cancer Discov 9, 1102–1123. 10.1158/2159-8290.CD-19-0094.

15. Plasko, G.R. (2021). The Functional Role of Tetranectin in Overnutrition-Induced Obesity and Hepatosteatosis in Female Mice. In T.U.o.T.H.S.C.a.S. Antonio, ed. Doctoral dissertation.

16. Kobayashi, H., Gieniec, K.A., Lannagan, T.R.M., Wang, T., Asai, N., Mizutani, Y., Iida, T., Ando, R., Thomas, E.M., Sakai, A., et al. (2022). The Origin and Contribution of Cancer-Associated Fibroblasts in Colorectal Carcinogenesis. Gastroenterology 162, 890–906. 10.1053/j.gastro.2021.11.037.

17. Xie, Z., Bailey, A., Kuleshov, M.V., Clarke, D.J.B., Evangelista, J.E., Jenkins, S.L., Lachmann, A., Wojciechowicz, M.L., Kropiwnicki, E., Jagodnik, K.M., et al. (2021). Gene Set Knowledge Discovery with Enrichr. Curr Protoc 1, e90. 10.1002/cpz1.90.

18. Buechler, M.B., Pradhan, R.N., Krishnamurty, A.T., Cox, C., Calviello, A.K., Wang, A.W., Yang, Y.A., Tam, L., Caothien, R., Roose-Girma, M., et al. (2021). Cross-tissue organization of the fibroblast lineage. Nature 593, 575–579. 10.1038/s41586-021-03549-5.

19. Han, H., Cho, J.W., Lee, S., Yun, A., Kim, H., Bae, D., Yang, S., Kim, C.Y., Lee, M., Kim, E., et al. (2018). TRRUST v2: an expanded reference database of human and mouse transcriptional regulatory interactions. Nucleic Acids Res 46, D380–D386. 10.1093/nar/gkx1013.

20. Zhang, X., Zhang, M., Sun, H., Wang, X., Wang, X., Sheng, W., and Xu, M. (2024). The role of transcription factors in the crosstalk between cancer-associated fibroblasts and tumor cells. J Adv Res. 10.1016/j.jare.2024.01.033.

21. Martinez-Zamudio, R.I., Roux, P.F., de Freitas, J., Robinson, L., Dore, G., Sun, B., Belenki, D., Milanovic, M., Herbig, U., Schmitt, C.A., et al. (2020). AP-1 imprints a reversible transcriptional programme of senescent cells. Nat Cell Biol 22, 842–855. 10.1038/s41556-020-0529-5.

22. Faure, L., Soldatov, R., Kharchenko, P.V., and Adameyko, I. (2023). scFates: a scalable python package for advanced pseudotime and bifurcation analysis from single-cell data. Bioinformatics 39. 10.1093/bioinformatics/btac746.

23. Dimitrov, D., Turei, D., Garrido-Rodriguez, M., Burmedi, P.L., Nagai, J.S., Boys, C., Ramirez Flores, R.O., Kim, H., Szalai, B., Costa, I.G., et al. (2022). Comparison of methods and resources for cell-cell communication inference from single-cell RNA-Seq data. Nat Commun 13, 3224. 10.1038/s41467-022-30755-0.

