## Supplementary figures and images for "Single-cell transcriptome analysis reveals evolutionarily conserved features during the transition from normal breast stromal cells to cancer-associated fibroblasts"

### Supplemental Figures

Supplementary Figure 1

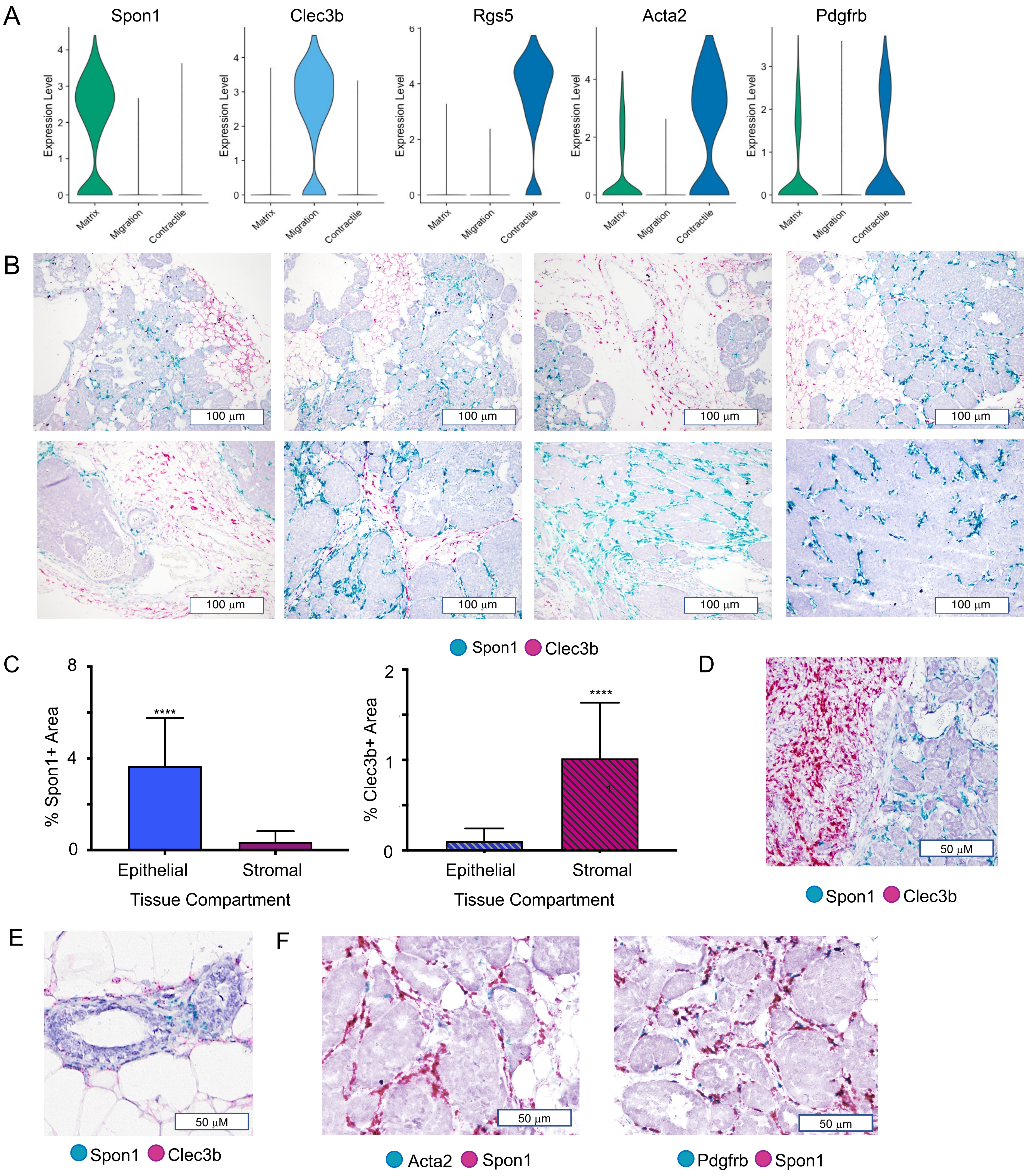

Supplementary Figure 2

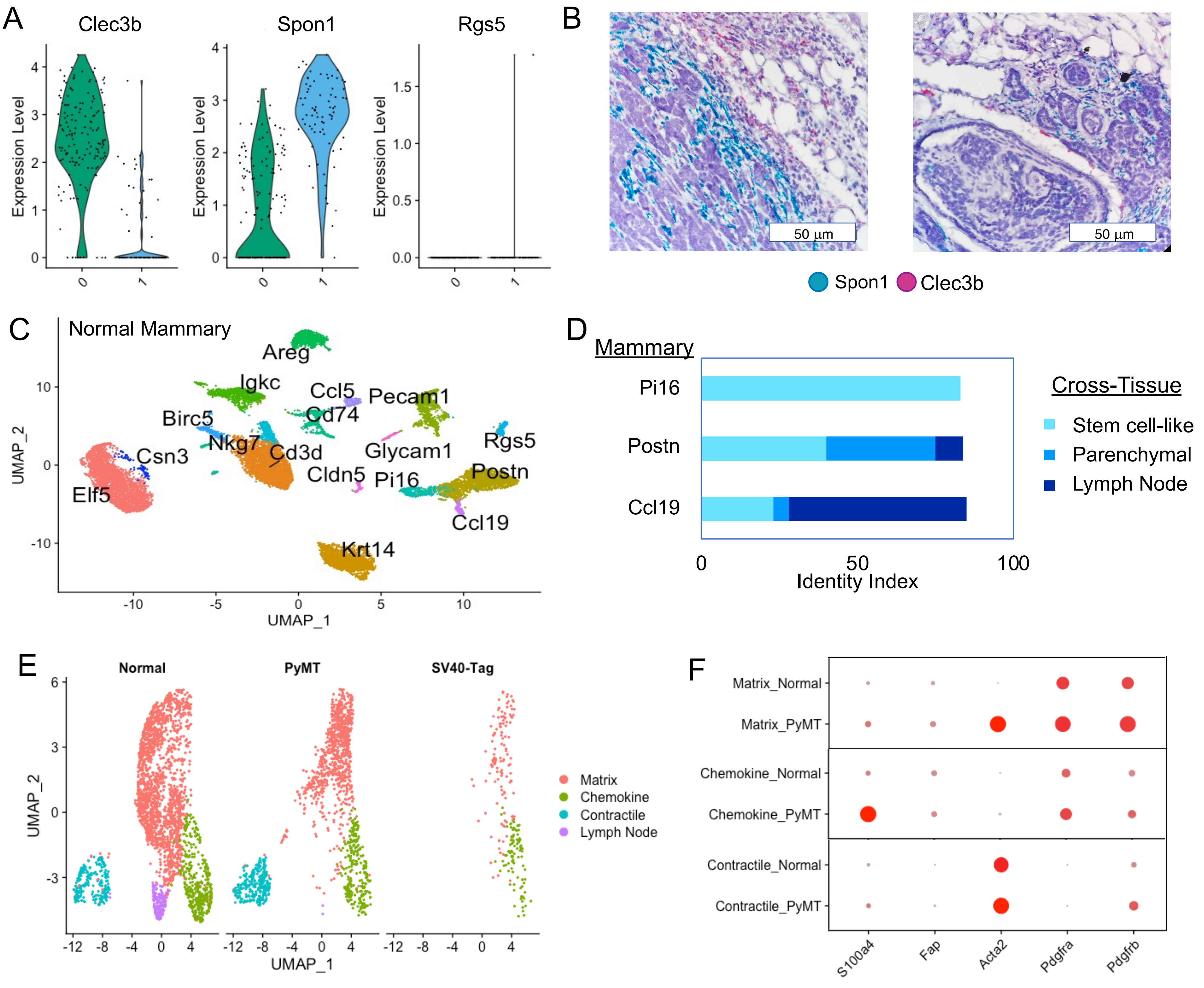

Supplementary Figure 3

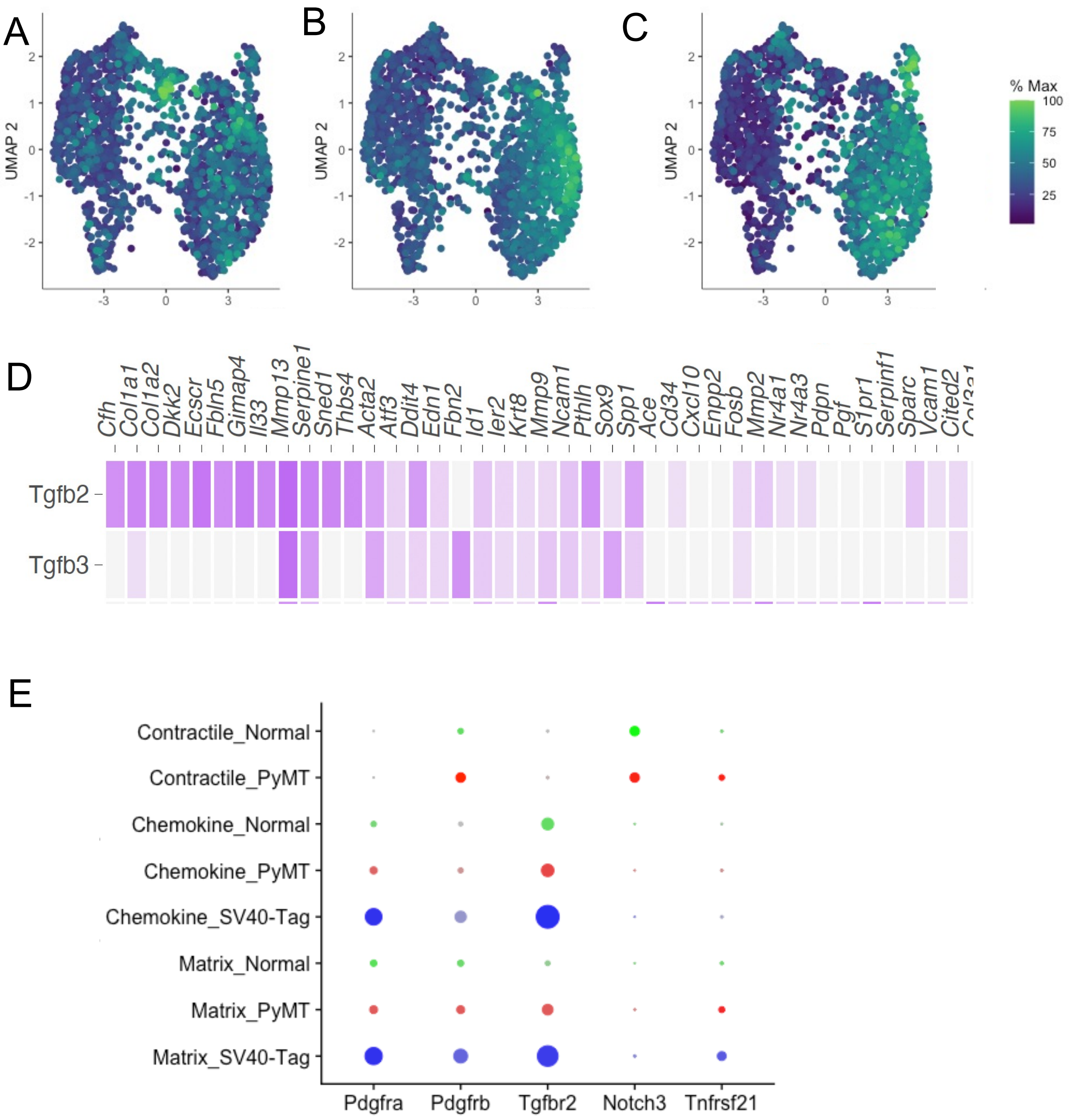

Supplementary Figure 4

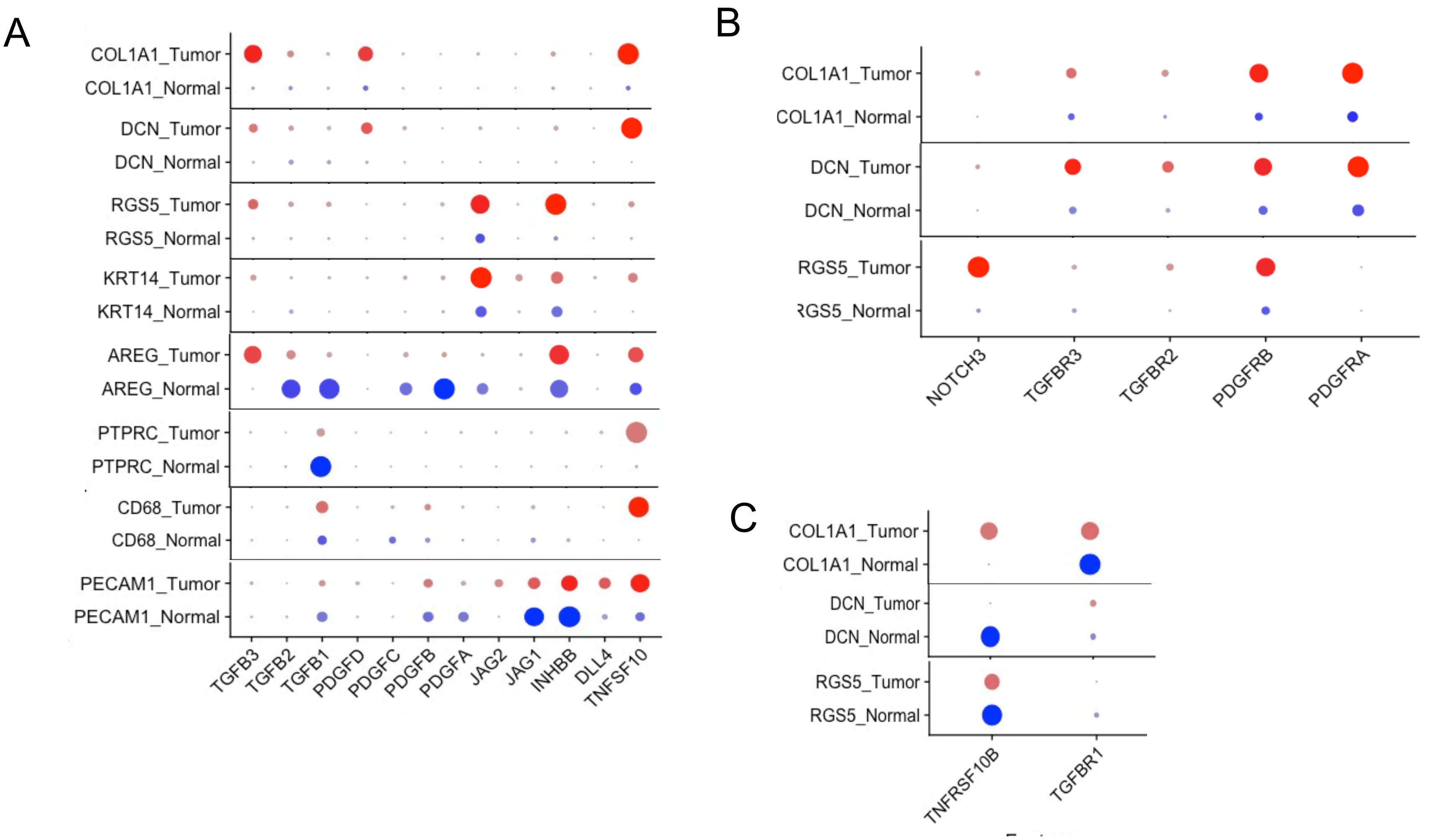
